# A pharmacokinetics-informed ODE extrapolates long-term fenofibrate transcriptomic responses

**DOI:** 10.64898/2026.08.25.746919

**Authors:** Yixuan Gao, Zheng Zhang, Yu Li, Jian’en Qiu

## Abstract

Long-term in vivo transcriptomic time courses are costly, limiting assessment of chronic molecular responses from short studies. We developed a pharmacokinetics-informed transcriptomic ordinary differential equation model (PKT-ODE) that links an oral pharmacokinetic profile and Hill drug-effect function to first-order turnover of co-expression modules. The model was fitted to rat liver responses to fenofibrate at three doses in Open TG-GATEs through day 8. At the held-out day-29 endpoint, PKT-ODE achieved Pearson *r* = 0.960 and mean squared error (MSE) = 0.148. In this dataset, these values achieved lower prediction error and higher correlation than four statistical baselines and validation-selected linear and multilayer-perceptron transition models. Literature-curated peroxisome proliferator-activated receptor α target genes occurred only in modules with positive fitted drug effects. These results provide a proof of concept for pharmacokinetics-informed transcriptomic extrapolation; cross-compound, cross-organ and alternative-regimen performance remain to be tested.

## Introduction

Drug-induced toxicity remains a major cause of late-stage candidate attrition ^1^, and molecular responses to repeated dosing can unfold over weeks. The temporal gap between an early molecular initiating event and a chronic apical outcome motivates adverse-outcome-pathway research and predictive toxicogenomics ^2–4^. Transcriptomic profiling resolves early responses at genome scale. Open TG-GATEs, DrugMatrix, the Comparative Toxicogenomics Database and the LINCS programme provide dose- and time-resolved expression resources ^5–9^. A central practical question is whether early transcriptomic responses can anticipate later molecular states. Reliable extrapolation could help prioritize chronic studies, although its effect on development timelines or animal use would require prospective evaluation.

Common extrapolation baselines carry the last observation forward or fit linear and log-time trends to early measurements. Learned transition models instead estimate a shared update from one molecular state to the next. In the implementations assessed here, elapsed time and administered dose are not explicit physical inputs. Pharmacokinetic–pharmacodynamic (PK–PD) models provide a complementary formulation. Indirect-response and turnover models describe delayed outputs driven by time-varying concentration, while the Hill equation maps concentration to a saturable effect ^10–16^. Applying this structure to coordinated transcriptomic states could make the assumptions about dose and time explicit.

Weighted gene co-expression network analysis (WGCNA) can summarize drug-responsive genes into modules. A first principal component then provides one directed score per module ^17–19^. We hypothesized that these scores could be approximated by first-order turnover towards a concentration-dependent state. This formulation assigns each module a baseline, signed drug effect and turnover rate, while dose and time enter through a pharmacokinetic driver.

Here we develop PKT-ODE and assess it as a proof of concept using the hepatic response to fenofibrate. Fenofibrate is a peroxisome proliferator-activated receptor α (PPARα) agonist with well-characterized effects on rodent lipid metabolism ^20–23^. We fit the model using data up to day 8 and evaluated the day-29 endpoint against statistical and learned comparators. We also tested whether fitted effect directions were concordant with known PPARα biology. The estimand throughout is the held-out day-29 module-condition mean for the same compound, organ, dose levels and daily regimen. This design evaluates within-compound temporal extrapolation, not generalization to unseen compounds, organs or dosing regimens.

## Results

### PKT-ODE couples pharmacokinetics to module turnover

PKT-ODE implements a three-stage computational chain from administered dose to transcriptomic state (Fig. 1). First, a one-compartment oral absorption model converts the daily dose *D* into the plasma concentration *C*_*p*_(*t*) of fenofibric acid. Once-daily repeat dosing is represented by linear superposition of single-dose profiles:

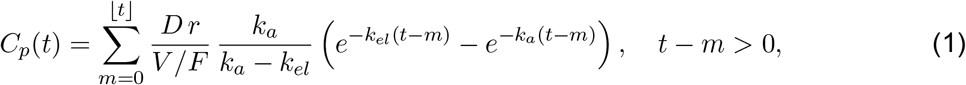

where *k*_*a*_ and *k*_*el*_ are the absorption and elimination rate constants, *V* /*F* is the apparent volume of distribution, and *r* converts parent dose to fenofibric-acid equivalents. Second, a Hill function maps concentration to a bounded drug-effect signal 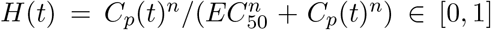. Third, each co-expression module *i* obeys a first-order turnover equation,

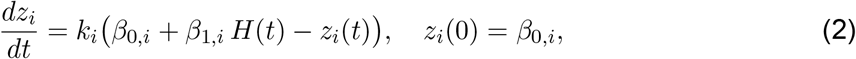

where *z*_*i*_(*t*) is the directed module PC1 score, *k*_*i*_ > 0 is its turnover rate, *β*_0,*i*_ is the fitted baseline offset and *β*_1,*i*_ is the signed drug effect. PK parameters were fixed from prior rat fenofibrate pharmacokinetics (*k*_*a*_ = 11.45 d^−1^, *k*_*el*_ = 2.64 d^−1^, *V* /*F* = 0.441 L kg^−1^, *r* = 0.884, *EC*_50_ = 5.42 μg mL^−1^ and *n* = 1). Only *k*_*i*_, *β*_0,*i*_ and *β*_1,*i*_ were estimated, giving 21 fitted parameters across seven modules. Each module was integrated by a semi-analytic scheme and fitted independently by bounded multi-start optimization (Methods). Under the fixed parameters, alternative doses or dosing intervals change *C*_*p*_(*t*) and therefore the predicted module states. Predictions under such alternative regimens were not evaluated here.

**Figure 1.**
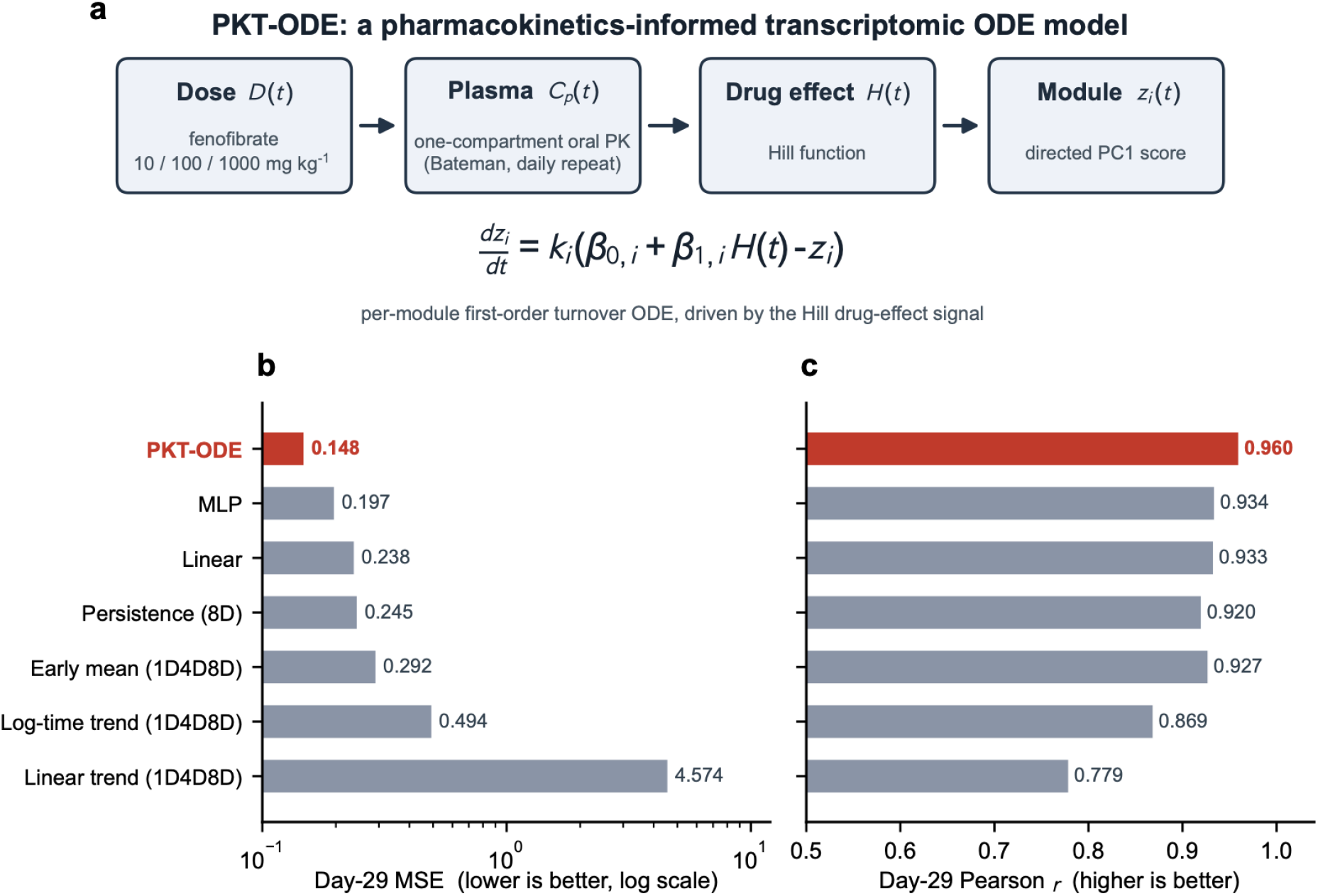
PKT-ODE overview and benchmark. **a**, The model couples administered dose *D*(*t*) to an oral pharmacokinetic profile *C*_*p*_(*t*), a Hill drug-effect signal *H*(*t*) and a first-order turnover ODE for each module *z*_*i*_(*t*). **b**,**c**, Held-out day-29 mean squared error (b, lower is better, log scale) and Pearson *r* (c, higher is better) for four statistical baselines, validation-selected linear and multilayer-perceptron transition models, and PKT-ODE (red). The statistical baselines were early mean, linear trend and log-time trend over days 1, 4 and 8, plus day-8 persistence. Metrics contain 21 module–dose values (three doses and seven modules).

### PKT-ODE extrapolates short-term data to the 29-day response

We fitted PKT-ODE using the first six sampling time points (3 h to day 8) across three fenofibrate doses (10, 100 and 1,000 mg kg^−1^; Table 1). Day 15 provided an intermediate held-out endpoint, and day 29 was reserved as the test endpoint (Fig. 2). At day 29, PKT-ODE achieved Pearson *r* = 0.960 and MSE = 0.148 across 21 module–dose condition means. At day 15, it achieved *r* = 0.971 and MSE = 0.154 (Table 2). A parameter-only re-simulation reproduced the day-29 correlation (*r* = 0.962), showing that the reported parameter values recover the numerical result (Methods). The fitted curves described the dose-dependent module states at the sampled times. These endpoint metrics are descriptive because the 21 values share doses, modules and the same underlying animals within each time point.

**Table 1.**
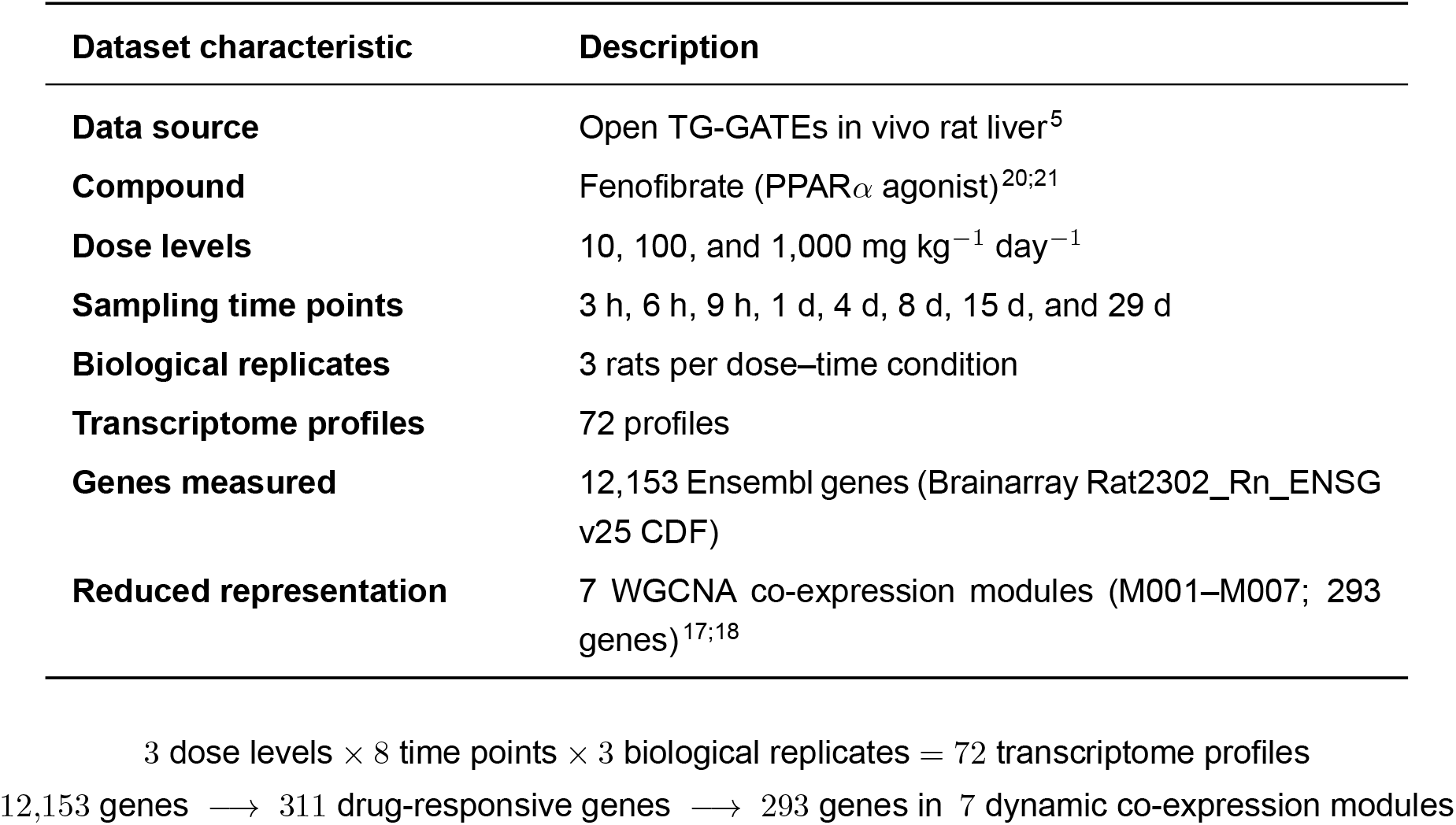
Dataset overview. Rat in vivo liver transcriptomic time course from Open TG-GATEs ^5^ used to fit and evaluate PKT-ODE. Once-daily oral fenofibrate was profiled at three doses and eight sacrifice time points with three biological replicates per condition; drug-responsive genes were summarized into seven signed WGCNA co-expression modules (M001–M007), whose directed first-principal-component scores served as the model state variables ^17;18^.

**Table 2.**
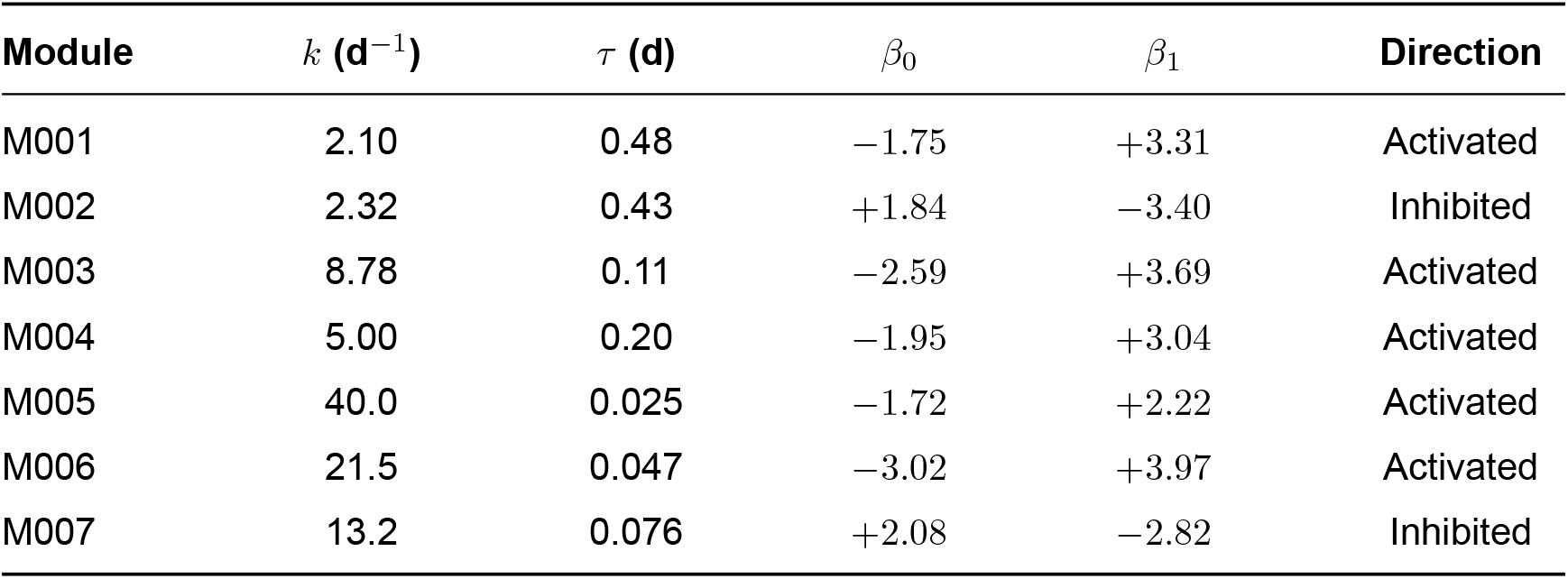
Fitted PKT-ODE parameters and per-split performance. Per-split performance across all modules and doses on the raw directed-PC1 scale: training (days 1–8), MSE = 0.164 and *r* = 0.897; validation (day 15), MSE = 0.154 and *r* = 0.971; and test (day 29), MSE = 0.148 and *r* = 0.960.

**Figure 2.**
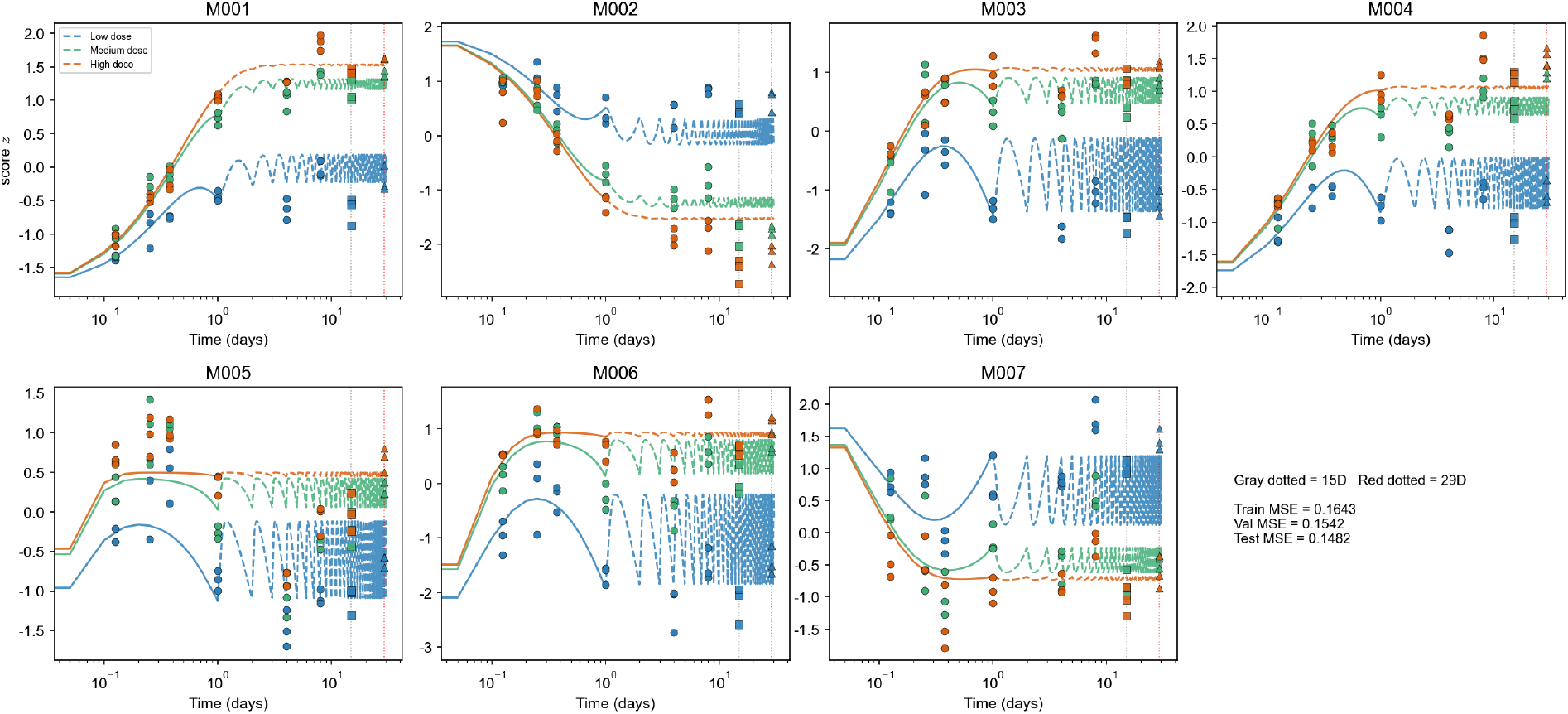
PKT-ODE fitted trajectories. Directed module PC1 scores for seven modules at low, middle and high dose (light-to-dark blue). Points show observed condition means and lines show fitted PKT-ODE trajectories from 3 h to 29 d on a logarithmic time axis. The shaded region beyond day 8 marks the held-out window, comprising the day-15 intermediate endpoint and day-29 test endpoint. Each panel reports the fitted turnover rate *k* and signed effect *β*_1_.

**Figure 3.**
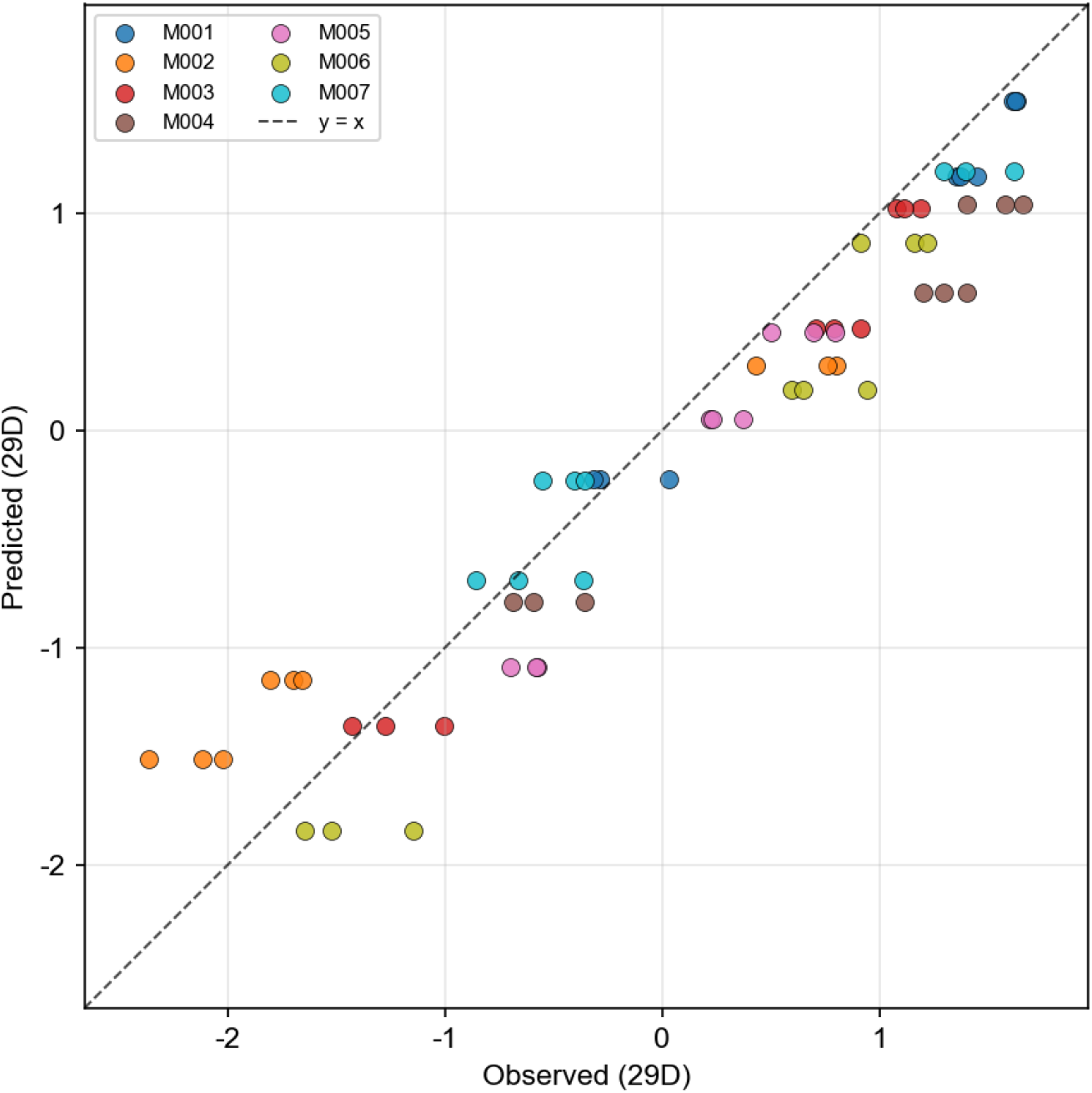
Day-29 predicted versus observed module scores. Each point represents one module–dose condition mean at the held-out day-29 endpoint and is coloured by module. The dashed line denotes identity. Across the 21 values, PKT-ODE achieved Pearson *r* = 0.960.

### PKT-ODE yields the lowest error among evaluated comparators

We compared PKT-ODE with four statistical baselines and two learned residual-transition models on the raw directed-PC1 scale (Fig. 1b,c). Learned-model checkpoints and configurations were selected by day-15 MSE before evaluation at day 29. PKT-ODE had the lowest day-29 MSE (= 0.148) and highest Pearson *r* (= 0.960) in this benchmark. The validation-selected multilayer perceptron achieved MSE = 0.197 and *r* = 0.934, while the validation-selected linear model achieved MSE = 0.238 and *r* = 0.933. PKT-ODE therefore had 27% lower MSE than the selected multilayer perceptron. Linear-trend extrapolation produced the largest error (MSE = 4.57, *r* = 0.779). Early mean (*r* = 0.927) and persistence (*r* = 0.920) retained high correlation but had larger errors. The learned models shared transition parameters across the three dose trajectories but did not receive dose or concentration as an explicit input. The comparison therefore supports the utility of PK-informed structure in this dataset, but it does not isolate which model assumption produced the difference.

### Fitted parameters indicate fast and slow transcriptional responses

The fitted turnover rates ranged from 2.1 to 40 d^−1^, corresponding to time constants *τ*_*i*_ = 1/*k*_*i*_ of 0.025–0.48 d (Table 2). M001 and M002 had the longest fitted time constants, whereas M005– M007 had values below 0.1 d. The M005 estimate reached the prespecified upper bound of 40 d^−1^ and should therefore be interpreted as boundary-limited rather than precisely estimated.

The signed effects classified five modules as increasing under the modeled drug signal and two as decreasing. These parameters provide a compact kinetic description, but their practical identifiability was not quantified.

### Modeled directions align with PPARα target genes

All seven modules were constructed from fenofibrate-responsive genes selected within the training window. We therefore assessed biological concordance by cross-referencing module membership with a 58-gene literature-curated PPARα target set ^22^. Twenty-seven targets occurred among the modeled genes, and all 27 were assigned to modules with positive fitted drug effects (Fig. 4). M001 contained 18 of these genes, including *Hadha, Hadhb, Hmgcl, Slc25a20, Ech1, Acot8* and *Cyp2j3*. The two modules with negative fitted effects contained none of the 58 targets. This distribution is consistent with known fenofibrate–PPARα regulation. It remains descriptive because the target set was literature-curated and the modules were derived from fenofibrate-responsive genes.

**Figure 4.**
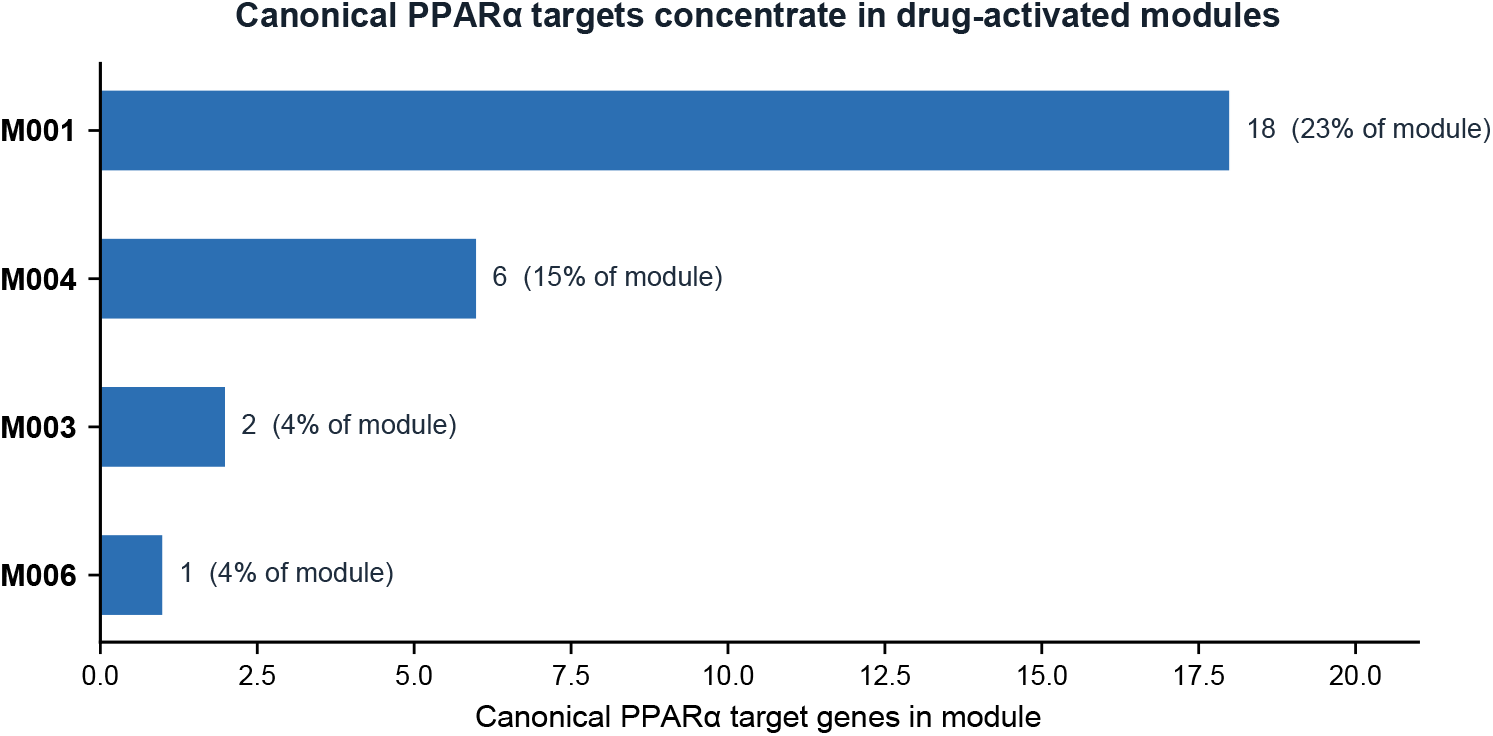
PPARα target-gene distribution across modules with positive fitted effects. Bars show the number of genes from a 58-gene literature-curated PPARα target set ^22^ in M001, M004, M003 and M006. Twenty-seven targets were present among the modeled genes, all in modules with positive fitted effects. The two modules with negative fitted effects contained no genes from this set and are not shown.

## Discussion

In this single-compound analysis, PKT-ODE extrapolated dose-dependent module states with lower day-29 error than the evaluated comparators. The same 21 fitted parameters described three doses because concentration entered the model explicitly. This structure also permits predictions at other times or regimens under fixed parameters, but that capability was not tested. Each module parameter has a direct mathematical role as a baseline, signed effect or turnover rate. Formal structural and practical identifiability analyses will be needed before these estimates can be interpreted as uniquely determined biological quantities.

The concentration-driven turnover formulation combines two established approaches. It uses the indirect-response structure and Hill concentration-effect mapping of PK–PD models ^10;11^, together with a low-dimensional co-expression representation from toxicogenomics. The concentration of literature-curated PPARα targets in modules with positive fitted effects supports biological concordance, but it is not independent validation of the model. In particular, the current analysis does not distinguish whether the benchmark performance arises from the PK driver, the smooth low-dimensional response, or both.

The potential toxicological value is narrower than direct toxicity prediction. PKT-ODE estimates later transcriptomic module states; it does not predict histopathology, clinical chemistry or an adverse outcome. The model could complement signature-based approaches by describing how a molecular response evolves over time ^2;4^. Establishing utility for safety decisions will require prospective evaluation across compounds and links to phenotypic toxicity endpoints.

Several limitations define the present evidence. First, the model was developed and evaluated on one compound in rat liver, with three animals per dose–time condition. Cross-compound, cross-organ and alternative-regimen performance is unknown. Second, day 29 is a single test endpoint, so denser chronic sampling is required to test the extrapolated trajectory. Third, the monotonic Hill driver cannot represent cumulative injury, immune feedback or non-monotonic responses. Fourth, fixed PK parameters were assembled from prior rat studies without uncertainty propagation or sensitivity analysis ^24;25^. Fifth, one turnover estimate reached its optimization bound, and no profile-likelihood or bootstrap analysis quantified parameter uncertainty. Finally, gene selection and module fitting used only the training window but remained specific to fenofibrate. The target-gene concordance is therefore descriptive rather than external validation.

Within these boundaries, PKT-ODE provides a proof of concept for linking pharmacokinetics to co-expression-module turnover. The current result supports within-fenofibrate temporal extrapolation and motivates broader validation before application to chronic toxicity prediction.

## Methods

### Data source and preprocessing

In vivo rat liver expression data were obtained from Open TG-GATEs ^5;6^. Raw Affymetrix CEL files were background-corrected, quantile-normalized and summarized by robust multi-array average using the Brainarray Rat2302_Rn_ENSG v25 custom CDF. Animals with any serious-pathology flag were excluded before normalization. Healthy controls had no pathology record and met all four biochemical thresholds: alanine aminotransferase ≤ 37.84 IU l^−1^, aspartate aminotransferase ≤ 81.72 IU l^−1^, blood urea nitrogen ≤ 16.92 mg dl^−1^ and creatinine ≤ 0.3684 mg dl^−1^. Log2 fold-change was calculated against the mean of at least three healthy controls matched by organ, sampling time and administration route. Fenofibrate was administered by daily oral gavage at 10, 100 and 1,000 mg kg^−1^. Liver expression was profiled at 3 h, 6 h, 9 h, 1 d, 4 d, 8 d, 15 d and 29 d, with three rats per condition.

### Co-expression modules and module scores

Gene selection and representation fitting used only the six time points through day 8; day-15 and day-29 values were excluded. A gene was selected when all three replicate signs agreed with the non-zero condition-median sign and the absolute median log2 fold-change was at least 0.5 in at least half of the 18 training conditions. This procedure retained 311 fenofibrate-responsive genes. Signed WGCNA with biweight midcorrelation used deep split 4, minimum module size 10 and merge cut height 0.05 ^17;18^. It yielded seven non-grey modules (M001–M007; 293 genes) and one grey set that was excluded. Fixed PC1 loadings, gene standardization and module-score standardization were fitted on the 54 training-window replicate profiles. Each PC1 sign was oriented to correlate positively with the mean standardized expression of its module ^19^. The fixed transformation was then applied to all 72 profiles. Replicate scores were averaged within condition, producing a 3 × 8 × 7 trajectory array (dose × time × module).

### PKT-ODE model

Plasma concentration of fenofibric acid after a single oral dose *D* followed a one-compartment absorption model (Bateman function) ^16^, and once-daily repeat dosing was modeled by linear superposition of single-dose profiles (equation in Results). The concentration was mapped to a bounded drug-effect signal by a Hill function ^14;15^ with coefficient *n* = 1 and *EC*_50_ = 5.42 μg mL^−1^. Each module obeyed a first-order turnover ODE with module-specific turnover rate *k*_*i*_, baseline *β*_0,*i*_ and signed drug effect *β*_1,*i*_, in the indirect-response tradition ^10–12^. PK parameters (*k*_*a*_ = 11.45 d^−1^, *k*_*el*_ = 2.64 d^−1^, *V* /*F* = 0.441 L kg^−1^, fenofibric-acid/parent mass ratio *r* = 0.884) were fixed from prior rat fenofibrate pharmacokinetics and were not estimated from the expression data ^24;25^.

### Integration and parameter estimation

Each module ODE was integrated on a 0.05-d grid over [0, 30.5] d. Within each interval, the driving term was linearly interpolated and the corresponding exponential update was evaluated analytically. Values at observation times were obtained by linear interpolation. The 21 free parameters (*k*_*i*_, *β*_0,*i*_ and *β*_1,*i*_ for seven modules) were estimated independently by minimizing residual sum of squares across the 18 training condition means. Optimization used L-BFGS-B with 12 starts, comprising six deterministic starts spanning slow and fast rates and both effect directions plus six pseudo-random starts. The lowest-residual solution was retained. Turnover rate was optimized on the log scale over [0.03, 40] d^−1^.

### Evaluation and baselines

Only time points through day 8 entered PKT-ODE fitting. Day 15 was withheld from PKT-ODE fitting and used to select the learned comparator configurations; day 29 was reserved as the test endpoint for all models. Performance was summarized by MSE and Pearson correlation after flattening the seven modules and three doses. These metrics were treated as descriptive because module–dose values are not independent biological replicates. Statistical comparators comprised the mean of days 1, 4 and 8; linear and log-time trends fitted to those days; and day-8 persistence. Learned comparators were linear and multilayer-perceptron residual-transition models initialized from day 1, trained against days 4 and 8, and early-stopped at day 15. For each learned family, the configuration with the lowest raw-scale day-15 MSE was selected before day-29 evaluation. All models were evaluated on the same raw directed-PC1 scale. A parameter-only re-simulation reproduced the reported day-29 correlation within 0.002 (*r* = 0.962 versus 0.960).

### PPARα target-gene cross-reference

Module genes were cross-referenced with a 58-gene PPARα target set curated from the cited literature ^22^. Ensembl identifiers were mapped to gene symbols using the Brainarray CDF description table. Target counts were summarized by module and compared descriptively with the sign of the fitted *β*_1,*i*_. No inferential test was applied to this target-gene distribution.

## Data availability

The raw transcriptomic data analysed in this study are publicly available from Open TG-GATEs (https://dbarchive.biosciencedbc.jp/en/open-tggates/desc.html). The database is provided by the Toxicogenomics Project and Toxicogenomics Informatics Project under CC Attribution-Share Alike 2.1 Japan.

## Code availability

The preprocessing and module-dynamics repository is available at https://github.com/ZhangZane-Westlake/PKT-ODE.

## Acknowledgements

This work was developed during the 2026 PEBBLE BioFusion Workshop at Westlake University in Hangzhou, China, from July 24th to August 4th.

## Author contributions

Yixuan Gao conceptualized the PKT-ODE model, designed its mathematical formulation and model architecture, implemented the model and training pipeline, performed model optimization and evaluation, and analysed the resulting transcriptomic trajectories.

Zheng Zhang conceived the overall study and formulated the central research question. He acquired and preprocessed the Open TG-GATEs transcriptomic data, performed quality control and feature preprocessing, implemented the dimensionality-reduction workflow, and designed and evaluated the baseline and conventional machine-learning models.

Yu Li established the compound-selection criteria, selected the compounds included in the study, and collected and curated compound-specific pharmacological, toxicological, and pharmacokinetic information.

Jian’en Qiu surveyed candidate dimensionality-reduction methods, designed and conducted comparative evaluations, assessed their robustness and biological interpretability, and contributed to the selection of the final dimensionality-reduction strategy.

Yixuan Gao and Zheng Zhang integrated the computational workflows and interpreted the modeling results. All authors discussed the results, contributed to the preparation and revision of the manuscript, and approved the final manuscript.

